# Optimization and Validation of Whole-Room Indirect Calorimetry for Improved Accuracy and Temporal Resolution

**DOI:** 10.64898/2026.04.10.717789

**Authors:** Christopher M.T. Hayden, Luke R. Arieta, Johanna M. Copeland, Michael A. Busa

## Abstract

With metabolic disease on the rise across the globe, the devices that can provide precise and reliable estimates of energy expenditure and macronutrient oxidation can play a critical role in the development and evaluation of therapeutic regimes and wearable technologies that can be used outside of the laboratory. Whereas, metabolic carts can provide short-term (minutes to hours) metabolic measurements, whole-room calorimeters enable long-duration (hours to days) metabolic assessment, providing insights into how metabolism changes in response to meals, activity, sleep, etc. Obtaining accurate metabolic measurement via whole-room calorimetry, however, requires rigorous methods for calibration and quality assurance. To date most room calorimeters have been tuned to assess energy expenditure over long periods of time, i.e. 24-hours. Here we present novel calibration and signal processing techniques and recommendations that aim to improve the utility of metabolic chambers for use over different measurement epochs. This work serves as both a transparent description of our hardware, validation procedures, and data processing approaches.

## 1 Introduction

Energy metabolism has been estimated through heat production – direct calorimetry – as long ago as the 1700s [7] with methods for measuring energy metabolism through gas exchange happen in 1862 [10] Accurately measuring heat production, however, is quite difficult and has considerable costs and limitations (described in detail in Kenny’s 2017 review [6]). Due to these constraints, researchers and clinicians shifted from directly measuring heat production to estimating heat production through the measurement of expired gases, a method termed indirect calorimetry. Indirect calorimetry has been validated against direct calorimetry and today is the basis for the majority of instruments used to measure whole-body energy metabolism [8].

### 1.1 What is indirect calorimetry?

In the human body, most energy is derived from the oxidation of carbohydrate, fat, and protein substrates through a series of processes collectively referred to as oxidative metabolism. During these processes, oxygen (O_2_) is consumed and carbon dioxide (CO_2_) is produced at rates directly related to the amount and type of substrate being oxidized. Indirect calorimetry is the measurement of the rates of O_2_ consumption *V̇*O_2_ and CO_2_ production *V̇*CO_2_ through the quantification of gas exchange between a subject and the environment. From *V̇*O_2_ and *V̇*CO_2_, estimates of energy expenditure (kilojoules/min) and substrate reliance (respiratory exchange ratio; RER) can be calculated. Common outcome variables from these estimates include total energy expenditure (TEE), sleeping or resting energy expenditure (SEE or REE), physical activity energy expenditure (PAEE), dietary-induced thermogenesis (DIT), and carbohydrate and fat oxidation rates. Additionally, while protein oxidation cannot be accurately determined from indirect calorimetry alone, it can be estimated if urinary nitrogen excretion is measured [14].

### 1.2 Types of indirect calorimetry

Of the many different modes of indirect calorimetry, the most popular and most widely used is the open circuit system, also referred to as flow-through respirometry. These systems come in the form of small portable units called metabolic carts, and full-size instrumented rooms called whole-room calorimeters (a.k.a. respiratory or metabolic chambers). Regardless of the mode or type of instrumentation, the purpose of all indirect calorimetry is to quantify the gas exchange - specifically *V̇*O_2_ and *V̇*CO_2_ - between a subject and the ambient air. From *V̇* O_2_ and *V̇* CO_2_ all other outcome variables are then calculated. Gas exchange is measured either through tubing connected to the mouth of a subject using a mask or ventilated hood or from the inflow and outflow air in a whole-room calorimeter.

While metabolic carts are far more common in research and clinical practice, the number of facilities with whole-room calorimeters is growing [2]. Most whole-room calorimeters resemble very small hotel rooms and are equipped with a bed, toilet, sink, computer, television, and exercise equipment. Whole-room calorimeters are useful tools for long-duration measurements because they do not require participants to wear masks or be attached via tubes and wires to any measurement device. This enables the analysis of energy expenditure in a pseudo-free living environment where subjects can eat, sleep, work, and exercise during measurement. The added utility of whole-room calorimeters over metabolic carts, however, comes with additional costs and limitations. Building, running, and maintaining a calorimeter is expensive and requires proper operation to ensure the accuracy of results.

### 1.3 Rationale and purpose

Whole-room calorimetry is an incredibly useful and increasingly more accessible technique for investigating human metabolism. To ensure the quality of metabolic measurement via whole-room calorimetry many steps must be taken to meet the general system requirements. Furthermore, to maximize the efficacy of whole-room calorimeters and minimize limitations, additional steps can be taken that enhance measurement accuracy and temporal resolution which serve to increase the amount data available from a data collection and open new opportunities to ask research questions across a range of measurement epochs.

The purpose of this manuscript is to describe and evaluate the performance of the whole-room calorimeters at the University of Massachusetts Amherst. This will include the introduction of new methods for assuring quality, improving accuracy, and increasing temporal resolution in a whole-room calorimeter. As most whole-room calorimeters are unique and custom-built, we will begin with an overview of whole-room calorimetry and a detailed description of the room and instrumentation at the University of Massachusetts Amherst. Subsequently, we will present data to describing how the calorimeter is calibrated and and data are processed; this will include both exisiting and novel methods for ensuring and improving the fidelity of whole-room calorimetric measurements. While the results reported here are specific to the UMass Amherst facility the methodologies that underlie the approach can be applied by others. The calibration, data processing, and quality assurance techniques will be applied as described here across all studies that are conducted at the site unless otherwise noted in the report of a specific study. We hope that the approach reported here can result in an avenue for whole-room calorimeter to be routinely used in an expanded set of questions in metabolic research.

### 1.4 Determination of *V̇* O2 and V*V̇* CO2 in whole-room calorimeters

As with all indirect calorimeters, the main purpose of a whole-room calorimeter is to measure *V̇* O_2_ and *V̇* CO_2_. Whole-room calorimeters act as large mixing chambers for gases, in which the expired air of the subject is diluted in ambient air. To calculate the contribution of the human subject to the gas concentration in a room, the volume of all inflowing and outflowing *O*_2_ and *CO*_2_ must be determined. Calculations can then be completed as follows:

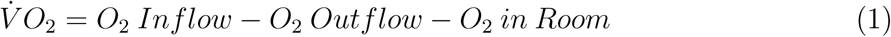

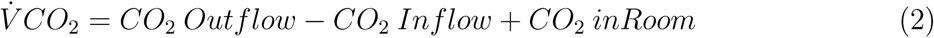

Equations 1 and 2 both contain three elements describing gas exchange *Inflow*, *Outflow*, and Δ *in Room*. The equations for *V̇* O_2_ and *V̇* CO_2_ are inverted because O_2_ is consumed whereas CO_2_ is produced. Because *O_2_ and CO_2_* are measured as a fraction or partial pressure of total volume, each part requires the measurement of both a volume (*Inflow Rate*, *Outflow Rate*, or *Room Volume*) and a concentration (fraction of O_2_ or CO_2_). In total, that makes six values that are needed for the calculation as shown for *V̇*O_2_:

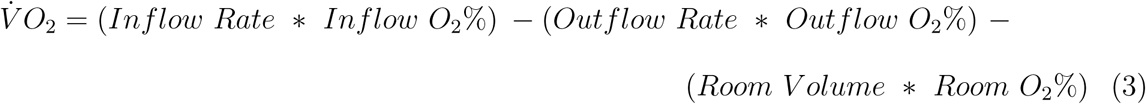

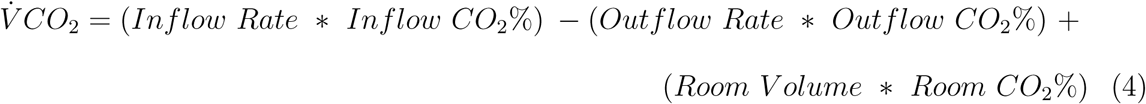

However, in practice only four of these terms (*Inflow Rate, Inflow gas%, Outflow gas%, and Room Volume*) are used to calculate *V̇ O*_2_ and *V̇ CO*_2_. Because room calorimeters are assumed to achieve perfect gas mixing, *Room O*_2_*%* and *Room CO2%* is equal to *Outflow O*_2_*%* and *Outflow CO*_2_*%*, respectively, thus in 3 and 4 *Outflow O*_2_*%* and *Outflow CO*_2_*%* are substituted for *Room O*_2_*%* and *Room CO*_2_*%*, respectively. Additionally - while very well sealed - the metabolic chamber is not completely airtight, the room is run at positive pressure (“push configuration”) causing any air leaks to be outward, and therefore only the inflow rate can be accurately measured. Due to this setup, *Outflow Rate* is estimated using the Haldane Transformation which assumes that there is no production or consumption of nitrogen by the person in the room. The Haldane Transformation [4], however, requires the addition of *Inflow CO_2_%* and *Outflow CO_2_%* values into the equation. Thus, the equations used in practice become much more complex (shown for *V̇O*_2_ in Figure 1).

**Figure 1:**
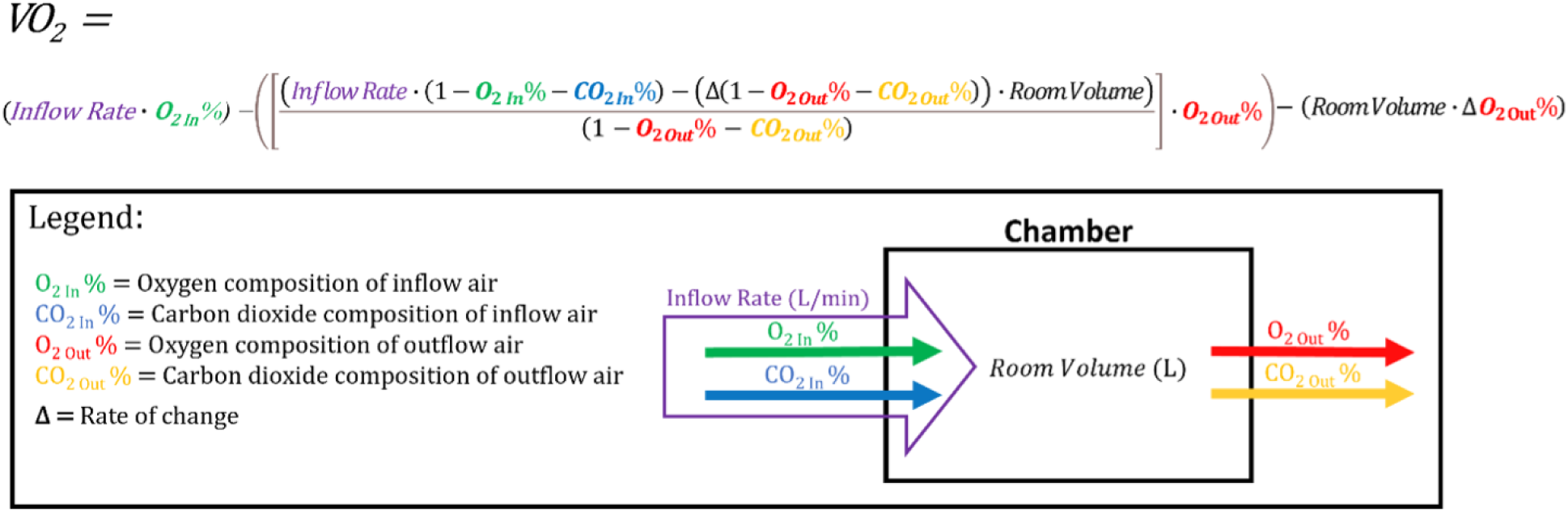
Equation for calculating oxygen consumption from a whole room calorimeter Given the potential for compounding error, it is clear that accurate and reliable determination of each of the six terms is required for high-fidelity measurement.

As seen in Figure 1., *Inflow CO_2_*% is used once, *Inflow Rate*, *Inflow O_2_*%, *Outflow CO_2_*%, and *Room Volume* are used twice, and *Outflow O_2_*% is used four times. Taking into account the common range for each measurement, it can be shown that under normal conditions the Δ*Outflow O_2_*% is the largest determinant of *V̇*O_2_ and, likewise, Δ*Outflow CO_2_*% of *V̇*CO_2_.

### 1.5 General requirements for whole-room calorimetry

#### 1.5.1 Environmental Control

Whole-room calorimeters require extensive control of experimental conditions to produce consistent and accurate results. To minimize gas exchange with unknown sources whole-room calorimeters must be airtight, including all windows, doors, and food/sample ports. Since no room is perfectly leakproof, pressure in the room must be kept positive to maintain the direction of any leak to be outwards. To maintain a positive pressure and a comfortable environment for the subject, the temperature must also be held relatively constant. Additionally, the air inflow rate must be tightly controlled as it impacts the pressure and temperature of the room, is utilized directly in the calculation of *V̇O*_2_ and *V̇CO*_2_ and is used to maintain *CO*_2_ at safe concentrations and within the analyzer measurement range. The composition of inflow air is also important, and as such is often sourced from medical-grade air filtration and air-drying systems with flow rates controlled using mass flow controllers (MFC). Outflow rates for the chambers can be controlled via MFCs as well and can be used to prevent pressure from getting too high or low in the chamber.

#### 1.5.2 Air Mixing

In a whole-room calorimeter, the gas that is expired by the subject immediately begins to mix with the ambient air in the room. Adequate gas mixing must occur to ensure that sampled gas is representative of the gas concentration of the entire room and not pockets of unmixed air. Powerful fans in the calorimeter are run continuously to maintain a well-mixed chamber and minimize airflow dead spaces such as those created by furniture.

#### 1.5.3 Precision Gas Analyzers

As room air mixes, gas samples exit through the sample port and travel to the gas analyzers in a separate room. Due to the large volume of the rooms, human subjects induce only small changes (∼thousandths to ten thousandths of a percent/min) in-room gas concentrations. Given the relatively small signal, measurement at this scale necessitates highly precise gas analyzers to maximize the signal-to-noise ratio. These analyzers are commonly custom-ordered to optimize the measurement range for calorimeters and have specific requirements for accurate measurement as follows. Sample humidity must be kept below specified levels to avoid condensation in the system and enable accurate gas concentration measurement. Fluctuations in humidity within the acceptable range must also be minimized using drying circuits or measured and corrected for. Sample flow rate must be held constant and within analyzer specifications, which can be completed using a combination of sample pumps and MFCs. To properly function, some oxygen analyzers also require a stable reference gas (typically from a precision mixed gas tank) to flow simultaneously over the sensors during measurement. Finally, gas analyzers must be calibrated regularly using mixed gases or a precision blender, and during data collection, gas concentrations must remain inside the analyzer’s measurement range.

### 1.6 Other considerations in whole-room calorimetry

When all requirements are met whole-room calorimeters can measure energy metabolism over long periods with high levels of accuracy, however, they are not without limitations. Room calorimeter experiments have high costs, both monetarily, and in the form of time commitment from the subjects and researchers, and they also have a relatively low temporal resolution compared to metabolic carts.

#### 1.6.1 Cost of Running Studies

Whole-room calorimeters are expensive instruments that require consistent maintenance to properly function, which can increase the cost to the housing facility. Experiments typically last for ∼23 hr and require staffing (generally researchers or nurses) for the entirety of the study. On top of staff wages, many facilities charge a fee for the use of the equipment to help cover the cost of the purchase and maintenance of the instrumentation. Running an experiment also has the added cost of feeding the subjects and compensating them for their time. Having consistent and reliable measurements can reduce the likelihood of needing to repeat or extend visits. For this reason, it is important to have comprehensive quality assurance that allows for testing, diagnosing, and resolving issues that may occur anywhere along the chain of measurement.

Maximizing the efficiency of data collection (i.e., the maximum amount of quality data in the least amount of time) is another means of keeping costs low and reducing the amount of time subjects need to stay in the room. Calorimeters often require subjects to be inside the chambers for an extended amount of time before data becomes “valid” (typically 30 minutes to several hours). This time can be decreased by bringing the gas concentrations of the room into optimal range faster by keeping the inflow rate low or by “priming” the room with N_2_ or CO_2_. Another potentially overlooked solution is fine-tuning analyzer calibrations to expand their optimal measurement range and improve their accuracy. Improving the accuracy of analyzers can also reduce the amount of time needed to calculate a given measurement such as REE, which is typically an average over a long period.

#### 1.6.2 Temporal Resolution

As described above, whole-room calorimeters have large volumes that cause a dilution of expired gas with ambient air. Imperfect mixing of gas can lead to a considerable temporal distortion in the outflow gas concentration signals. This distortion contributes noise to the signal, further decreasing the signal-to-noise ratio. To combat this noise and account for longer mixing time, the rate of change of gas concentrations (Δ*Outflow O_2_*% and Δ*Outflow CO_2_*%) is calculated using multi-point centered derivatives much larger than the sample rate. This derivative (i.e., rate of change) term, is often several minutes in length and while it helps to reduce noise and smooth outputs, it also causes a temporal broadening of events and a reduced event amplitude. This reduces the efficacy of using a whole-room calorimeter to evaluate events of short duration and transitions between events. Any effort that decreases noise and increases the temporal resolution of the instruments thus has the potential to increase their efficacy for answering research questions commonly reserved for metabolic carts. Many groups have devised complex post-processing methods including wavelet transforms [1, 11], as well as various filtering and spline fitting [2, 5, 3, 9, 13], however, these approaches can be overcomplicated and run the risk of removing signal along with noise. To decrease the need for such data transformations it is beneficial to decrease the noise in the output signal first at the measurement system before completing post hoc adjustments.

## 2 Description of Facility

### 2.1 Whole-room calorimeters at the Institute for Applied Life Sciences

The Center for Human Health and Performance in the Institute for Applied Life Sciences houses two whole-room calorimeters, one large (4.0 x 3.4 x 2.6 m) and one small (1.2 x 2.4 x 2.4 m) which were custom built in-place (MEI Research, Ltd, Edina, Minnesota). The larger calorimeter contains a bed, sink, toilet, treadmill, TV, desk, window, and two pass-through ports, with ample free space in the middle of the room (Fig. 2 A). This room is larger than most (32,500 L) and was designed to enhance participant comfort and allow for analysis of various activities of daily living that require moderate ambulation. The smaller room or “flex” calorimeter was designed to evaluate metabolism during stationary exercise or rest, with its lower volume (6,500 L) providing better accuracy and finer temporal resolution. The flex calorimeter is versatile as it can house equipment such as a stationary bike or be fitted with a bed reducing its volume to 5,000L - See Figure 2.

**Figure 2:**
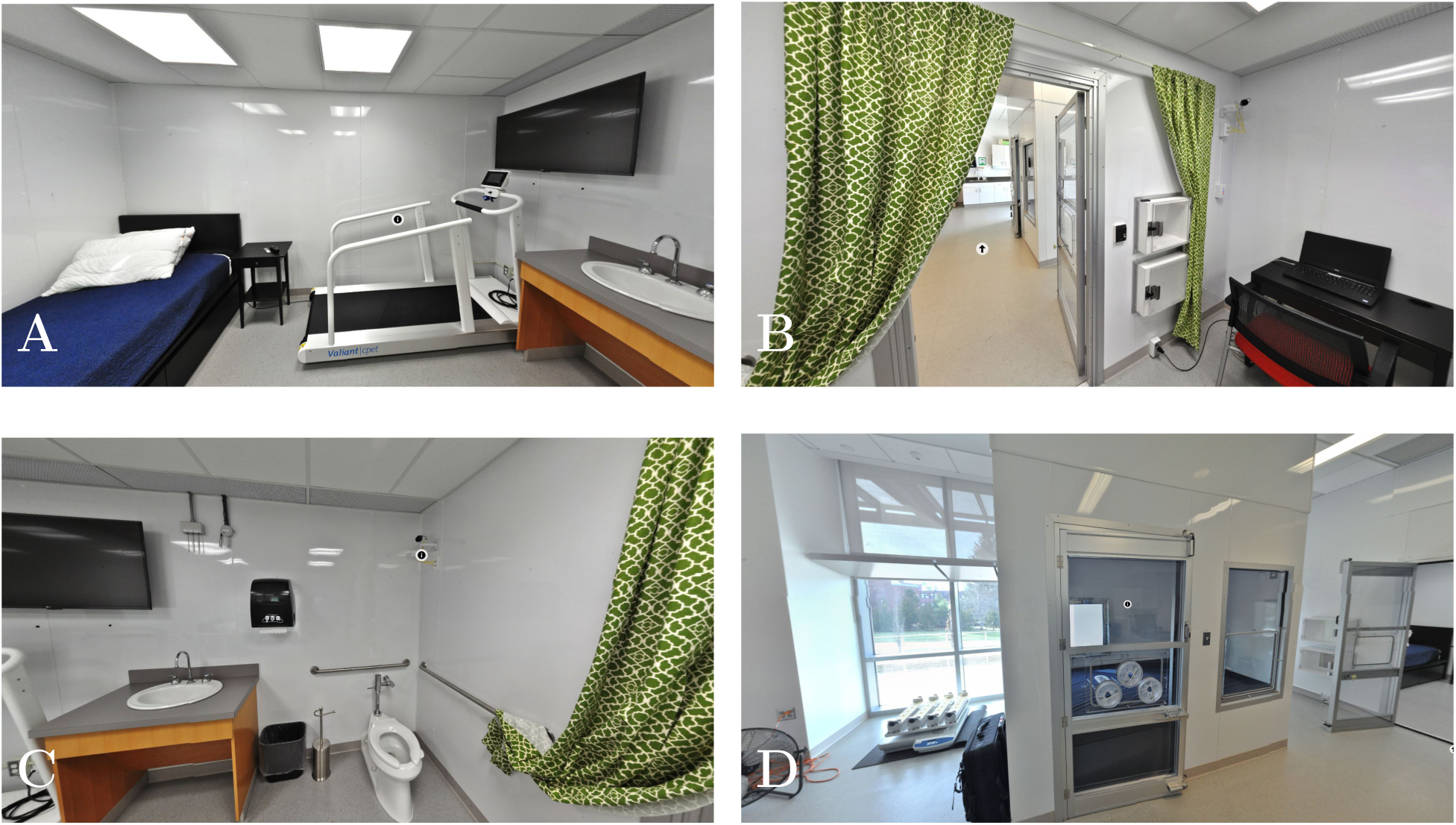
**A:** Bed & Treadmill Large (32,500L) Chamber; **B:**Ports and Door Large Chamber; **C:**Sink and Toilet Large Chamber; **D:** Exterior View of Small (5,000L) Chamber

### 2.2 Airflow Circuits

Inflow air for each chamber is provided by a medical air filtration system in a lower level of the building, this ensures air is free of airborne particles and is sufficiently dry (dew point < −30° C) as it is also used as a purge gas in the drying circuit. The air is then piped up into the equipment room where sample lines remove a small amount of air that then flows to the inflow analyzer (Fig. 3). Next, the air is passed through mass flow controllers and into the corresponding calorimeters where it is circulated by in-unit fans. The air is then pushed out by positive pressure, exiting through the outflow circuit piping. Sample air lines remove air from the calorimeter via sample pumps (Air Dimensions, Inc., Deerfield Beach, Florida) and flow it to one of two outflow analyzers – one for each room. Since sample air can regain humidity from the room and participant, outflow gas samples are dried between the room and outflow analyzers by a sample dryer (PD-Series, Perma Pure LLC, Lakewood, New Jersey) to ensure that fluctuations do not interfere with measurements and to prevent condensation. As an assurance measure, sample H_2_O sensors (Measurement Specialties Inc, Hampton, Virginia) sit before each analyzer to verify adequate drying.

**Figure 3:**
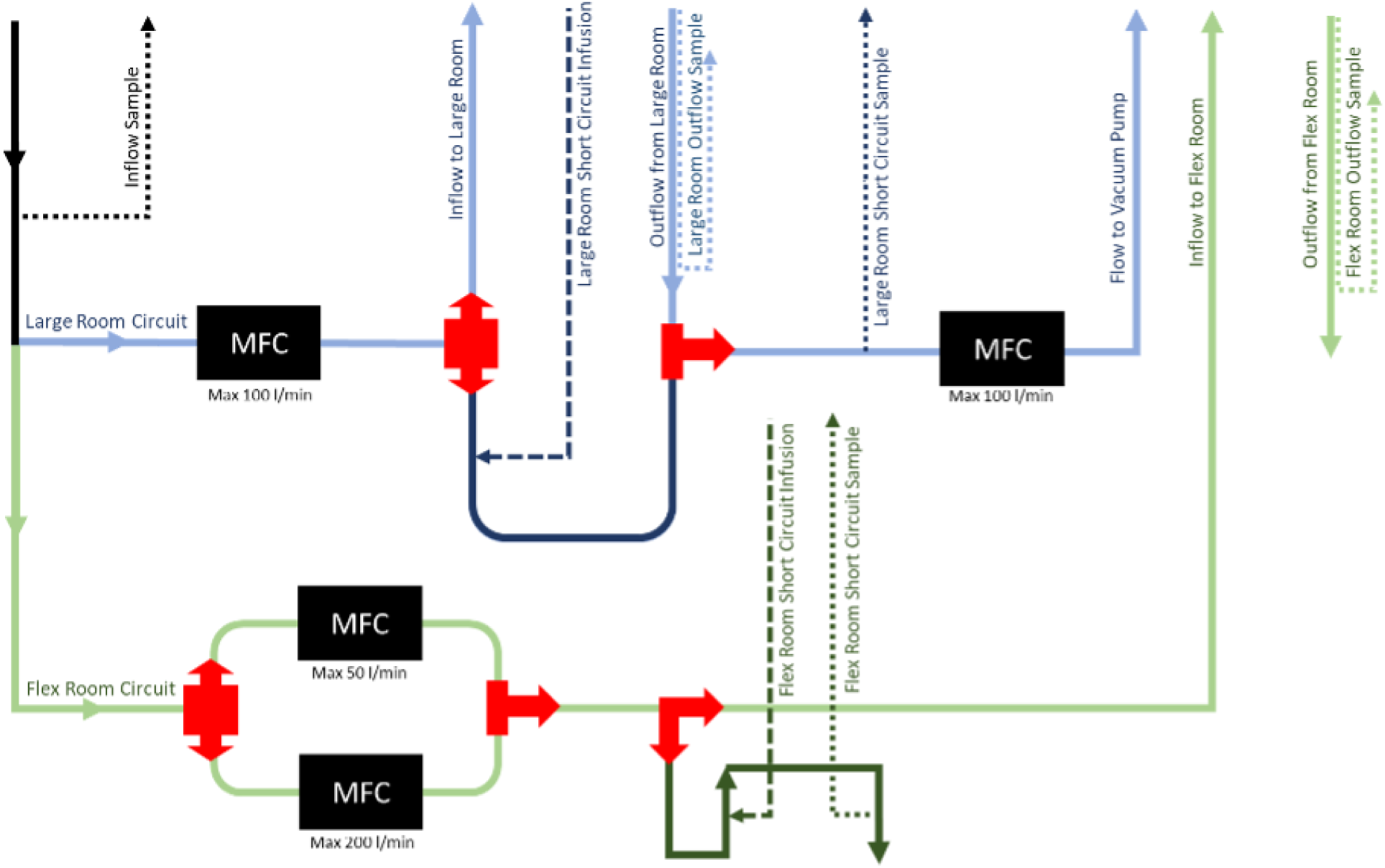
Calorimeter airflow circuits. Schematic of the airflow to and from the large and flex room calorimeters, the sample and infusion lines, and the short circuit configurations. MFC, mass flow controller.

A push calorimeter is a system in which air is actively forced into the chamber, creating an internal positive pressure, and causing all leaks to flow outward. The advantage of this system is that the inflow air comes from only one source and can therefore be easily controlled (making it highly stable) and it does not require the construction of a buffer space or other means of controlling the air composition surrounding the chamber. It also allows for the use of blood sampling ports (Fig. 2D) without the risk of contaminated air being pulled in. The disadvantages of a push system are that small but likely unavoidable outward leaks cannot be measured. Due to this, gas mixing in the chamber must be highly effective so that any given leak can be assumed to have the same composition as the output air and therefore does not cause error in measurement [12].

Our large calorimeter functions as a push system with inflow air controlled by an MFC (Porter, Parker Hannifin Corporation, Cleveland, Ohio) with a max flow of 100 l/min. The flex calorimeter is also a push chamber and has two inflow MFCs - max flows of 50 and 200 l/min - allowing for accurate flow control at both very low (resting) and high (exercise) metabolic rates. The large calorimeter has the added utility of optionally running in a “push-pull” configuration by utilizing a vacuum blower and an MFC on the outflow circuit. This configuration enables operation at a lower pressure to minimize leaks, which is especially useful when high inflow rates are needed (i.e., intense exercise). Data discussed in this manuscript were collected in push-only configuration.

In addition to the standard airflow circuit described above, each of our calorimeters is equipped with a “short-circuit”. In this configuration, medical air bypasses the chamber and enters a short-circuit plumbing loop (dark blue in Fig. 3). Inflow air is sampled as before, but outflow air is now sampled from the plumbing prior to the MFC (Fig. 3), instead of the room itself. Additionally, blended gases can be infused into the short-circuit air in between the inflow and outflow analyzers allowing the ability to test the measurement equipment without the potentially confounding factor of flowing gas into the chamber and sampling from it. This allows for tests with a chamber of effectively zero volume and is important for evaluating hysteresis in measurements and eliminating the mixing effects of gas in the rooms.

### 2.3 Analyzers

All gas samples are analyzed by one of three dual measurement gas analyzers (ULTRA-MAT6/OXYMAT6, Siemens, Munich, Germany), of which one analyzer is designated for analysis of inflow gas and the other two for outflow from each calorimeter. Each analyzer is equipped with both a paramagnetic O_2_ sensor and an infrared CO_2_ sensor. The analyzers provide a digital data signal representing the percent gas concentrations of air volume. The O_2_ analyzer requires a flowing reference source, which is provided by a custom mixed gas cylinder (∼21% O_2_, balance N_2_) with a two-stage regulator. These analyzers can accurately measure at flow rates of 0.3-1 l/min O_2_ and 0.3-1.5 l/min CO_2_, however, this accuracy only holds for the flow rate at which the analyzers were calibrated. Transient fluctuations in flow rate, therefore, have the potential to impart measurement error that is nearly impossible to correct *post hoc*. At a minimum flow rate must be controlled by rotameters, however, our lab utilizes MFCs (Alicat Scientific, Inc., Tucson, Arizona) for enhanced control. All three analyzers can sample from inflow air and calibration sources, allowing for measurement comparison and quality control, the outflow analyzers can also sample air from the calorimeter rooms and the short-circuit. Because the sensors must be warmed up before measurements become accurate, air is constantly supplied to them between studies. The sensors in gas analyzers are also sensitive to vibration so care was taken to physically isolate them from other equipment. For example, sample pumps are connected to gas analyzers with flexible tubing instead of stainless steel to minimize the transmission of vibrations.

### 2.4 Precision Gas Blending System

To calibrate, validate, and perform modeling experiments, our room calorimeter facility is equipped with a precision gas blending system (MEI Research, Ltd, Edina, MN, USA). This system consists of 4 Mass Flow Controllers controlled with CalRQ software (MEI Research, Ltd, Edina, Minnesota) that allow us to combine pure O_2_, CO_2_, and N_2_ at individual rates from 0.015 to 20 l/min. Precision blended gasses can be directly sent to the analyzers, into either room or into either short-circuit. Gas tanks include pre-blended zero (21% O_2_, balance N_2_) and span (20% O_2_, 1% CO_2_, balance N_2_) tanks as well as tanks of pure O_2_, CO_2_, and N_2_ gas.

### 2.5 Environmental Control

Each calorimeter is equipped with temperature, humidity, and relative pressure sensors. Chamber temperature is tightly controlled (<0.05°C change from setpoint per minute) over a wide range of setpoints (15-35°C) through a custom-built HVAC system (Harris Environmental Systems, Andover, Massachusetts) with a programmable logic controller. Such tight temperature control allows us to ensure that any changes in room pressure are slow. Additionally, inflow and outflow rates are controlled by proportional-integral-derivative (PID) control via CalRQ software. This PID control of inflow rate maintains CO_2_ concentration to safe levels (target < 0.3% CO_2_) while the outflow PID control ensures that the chamber maintains adequate positive pressure (when running in push-pull configuration). If desired, PID control can be disabled and flow rates can be manually set, this often is used during calibration procedures.

### 2.6 Participant Activity Monitoring & Interaction

Participant activity in the chamber can be monitored in several ways. Indirectly, motion can be identified via microwave motion sensors (Museum Technology Source, Inc., North Reading, Massachusetts). Direct observation can be made via plexiglass viewing windows into rooms and voice communication via handheld 2-way radio or intercom. Additionally, the large chamber is outfitted with two time-synchronized video cameras (Noldus Viso, Wageningen, NL), which allows for the contextualization (e.g. sleep, wake, exercise, ADL) of metabolic measures during data analysis. Importantly, participants can suspend recording intermittently from inside the chamber by the push of a button affording them privacy while changing clothes or using the toilet. Additionally, the cameras are positioned such that they do not view the toilet. The large chamber contains two pass-through box ports for the exchange of meals and samples between researchers and participants. Custom ports can also be installed on either calorimeter door to facilitate blood draws while maintaining a near-constant seal on the door allowing for minimal data loss (See Fig. 2A-C).

### 2.7 Third-Party Device Integration

Each chamber is equipped with a USB and an RS-232 serial port that allows for additional instrumentation to be used within the chamber. Our facility houses both a treadmill (Lode, Valiant 2, Lode. B.V., Groningen, NL) and a Velotron cycle ergometer (Sram Inc, Chicago, USA). From the CalRQ software, profiles can be applied that specify treadmill speed and grade and cycle wattage or resistance. Live measurements from these devices are integrated with the metabolic measures. The treadmill is not placed in the small chamber for safety reasons. Other devices can be integrated so long as drivers can be integrated with CalRQ. Additionally, the system time for the computer that operates the chambers is provisioned from our local Network Time Protocol server (Masterclock Inc., St. Charles, MO, USA) which allows for the synchronization of other relevant systems within the UMass Center for Human Health & Performance (e.g., wearable devices, video, and other nonendemic devices).

## 3 Chamber Validity

To ensure that whole-room calorimeter data accurately and reliably represents the metabolism of test subjects it is necessary to 1) validate the system, and 2) continually ensure the system remains valid (i.e., quality assurance). Because whole-room calorimeters consist of equipment from multiple manufacturers and are variably constructed to fit into existing infrastructure validation can be difficult. While each component may meet individual manufacturer specifications and tolerances when they are combined new issues may arise that can impact measurements. As a result, having the ability to test reduced sets of components can aid in validation and troubleshooting if issues arise. These subsystems can then be combined to evaluate total system performance. Here we describe procedures that can be used to ensure the calibration, validation, and quality assurance of metabolic chambers and pinpoint sources of error throughout the various instrumentation.

### 3.1 Mass Flow Controllers - Metrology

The precision blending system and inflow MFCs are calibrated at MEI Research’s traceable metrology lab semi-annually. The metrology lab is comprised of two displacement flow calibrators: one 5-50,000 ml/min (ML800, Mesa Labs, Lakewood, CO) and the other 5 −500 l/min (ML1020, Mesa Labs, Lakewood, CO). These calibrators undergo recalibration by Mesa Labs annually. Each MFC calibration includes 10-point tests across the full-scale range (10 −100%, 10% steps) for each applicable gas (air, O_2_, CO_2_, N_2_). A linear 10-point correction curve is applied to the MFC for each applicable gas, *k* factors are not used as this method has proven to make a small but meaningful difference. Once the calibration curve is applied, a second 10-point test is run to ensure the error in the MFC data is less than 0.5% across 5-95% of the operating range. If the device does meet this criterion, it is inspected for damage and if no damage is found, the full calibration procedure is repeated.

### 3.2 Gas Analyzers

The Siemens gas analyzers can be calibrated directly on the instrumentation panel using pre-mixed gas cylinders with known concentrations of O_2_, CO_2_, and N_2_ that span the operating range of the analyzers: zero (21% O_2_, balance N_2_) and span (20% O_2_, 1% CO_2_, balance N_2_). Each gas is run across the O_2_ and CO_2_ sensor for each analyzer for ∼10 min. After readings have leveled out the measured values are reset to match the known values. This is referred to as a *hard calibration*, as it resets the correction factors within the equipment, changing the raw values recorded by the software, and cannot be undone. Hard calibrations are performed anytime there are adjustments to the system that may affect sensor measurement such as changes in sample flow rate or reference gas concentration and are also advised anytime the analyzers are moved or displaced (e.g., during maintenance). Recordings during calibration are stored in case troubleshooting or retrospective review is required.

### 3.3 Precision Gas Blending

A precision gas blender, made up of multiple MFC’s, can be used to test the performance of the chamber. The system mixes pure gases (from O_2_, CO_2_, and N_2_ tanks). These can be mixed in ratios that simulate metabolic rates, or test specific performance metrics. The blended gases are mixed with the *inflow* air after the inflow analyzer and these mixture of these gasses is then measured by the *outflow* analyzer; because the *infused* gasses are measured the precision of the entire hardware and software system can be evaluated and tuned.

This technique has several advantages over propane combustion procedures that are commonly used for validation [2]. Gas infusions are not limited to simulating a single metabolic rate, allow for RER in a physiological range for humans, and can utilize premade profiles of different *V̇*O_2_, *V̇*CO_2_, and RER values. These advantages make it possible to test chamber reactivity to various simulated physical activity protocols. Further, because the gas profile time series are known with a high level of precision, the expected output measurements can be determined at the sample level and the effects of post-processing decisions can be evaluated. Additionally, while not covered here, infusion rates can be changed in a precise and near-instantaneous manner, to identify chamber parameters such as reaction time or minimum detectable difference.

#### 3.3.1 Short Circuits

In practice, running short-circuits first enables the evaluation of accuracy, reliability, and reactivity of the measurement equipment independent of flowing gas through the chambers. This has the advantage of allowing the operator to eliminate sources of error in the complex chamber system. For example, an issue with air leaks in the plumbing. To go one step further, if a problem with the gas analyzers is suspected, blended gas can be flown directly to the sensors circumventing inflow air and any additional plumbing. The ability to repeatedly provide the same input (blended gas infusions) to segmented portions of the calorimeter system enhances traceability over time, which is critical to allow for data interoperability over time as components of the system fail, are changed, or added to the system. For this reason, infusion tests (room, short-circuit, and direct to analyzer) should be run regularly – either prior to a new protocol or at a minimum every 3 months - to provide data for quality assurance, troubleshooting, and identifying targets for future system improvement.

#### 3.3.2 Infusions

Similar to *short circuits*, *infusions* infuse known mixes of *CO*_2_ and *N*_2_ into the metabolic chamber. The purpose of this is to move beyond just evaluating measurement precision of the gas flow and analyzer systems and move to understanding how the chamber might response to e.g., different metabolic rates, temperatures, sampling rates, etc. Leveraging this capacity has allowed us to conduct testing that provides us objective information from which we can make operational decisions that impact study design, data sampling, and signal processing.

## 4 Chamber Reliability

### 4.1 Calibration Checks

*Calibration checks* are performed weekly and before studies by flowing the same zero and span gasses directly over the analyzer and verifying that measured values are within a tolerance range (± 2% error). Because consistency in measurement across the analyzer range is more important than the absolute value, the difference between expected and measured gas readings is used to calculate percent error as shown below for O_2_.

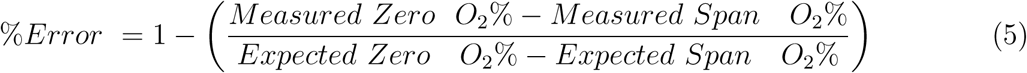

Data streams are also recorded during calibration checks and calculated error results are logged to improve traceability over time. The same process is followed for the CO_2_ sensor.

### 4.2 Null tests

After calibration checks are passed *null tests* are conducted as another measure of quality control. This consists of running medical air directly over both inflow and outflow analyzers, during which time the gas concentrations from both analyzers are nearly the same, and *V̇* O_2_ and *V̇* CO_2_ are approximately zero. Null tests are performed at the start and end of every data collection and are used to check for analyzer drift. While inflow air concentrations from pre to post-test may change, any offset between analyzers should not. Null tests can also be performed as a stand-alone diagnostic check to ensure that all analyzers are producing accurate readings and are beneficial for longer evaluations because they don’t consume pure or mixed gas from tanks.

### 4.3 Short Circuits & Infusions

As described in sections Short Circuits and Infusions these processes can be used periodically to track the stability of chamber performance in response to wear and tear, environmental changes, and changes to the larger infrastructure that support the buildings the chambers are located. Additionally, when new protocols are conceived that might go beyond the ranges that the chamber has previously or routinely tested at. The data from these simulations can used to evaluate things like required sampling rates, derivative needed to gain the appropriate temporal resolution - while balancing signal/noise, the amount of time you might need a person in the chamber before they equilibrate.

## 5 Chamber Performance

With the stability and data quality afforded by the practices described above, further efforts to enhance chamber performance can be focused on data analysis strategies. Here we evaluate the effects of various post-processing methods on measurement accuracy and temporal resolution. The goal of this is to improve data collection efficiency, identify sources of potential error in human data where the expected values are unknown, and enable accurate measurement over short epochs and during periods with high variability in metabolic rate.

### 5.1 Infusion Protocol

To validate our measurements and test the effects of different signal processing methods we completed a series of gas infusions. Specifically, we completed 4-hr N_2_ and CO_2_ infusions to simulate *V̇* O_2_ and *V̇* CO_2_ during 2 hr of steady-state (“flat” portion), eight 5-minute step changes in metabolic rate (“dynamic” portion), and 30 min of a simulated treadmill walk (Fig. 4). These “simulation infusions” were completed in duplicate at estimated metabolic equivalents for subjects with body weights of 30, 50, 70, 90, 110, 130, 150, and 170 kg, making sixteen total infusions. The first hour of each infusion was discarded to allow the room to equilibrate and allow for signal quieting. Inflow rate was held constant at 70 L/min during the infusions and all measurements were sampled at 1 Hz. All infusions were completed in our 32,500 L metabolic chamber - which poses a more difficult condition than the 6,500 L chamber for accuracy and temporal resolution measurement due to the larger volume and longer air turnover rate. We calculated the area under the curve error (AUC, %) as the difference between the expected vs. the measured values during different periods of interest within this infusion to evaluate the effect of post-processing strategies on measurement accuracy.

**Figure 4:**
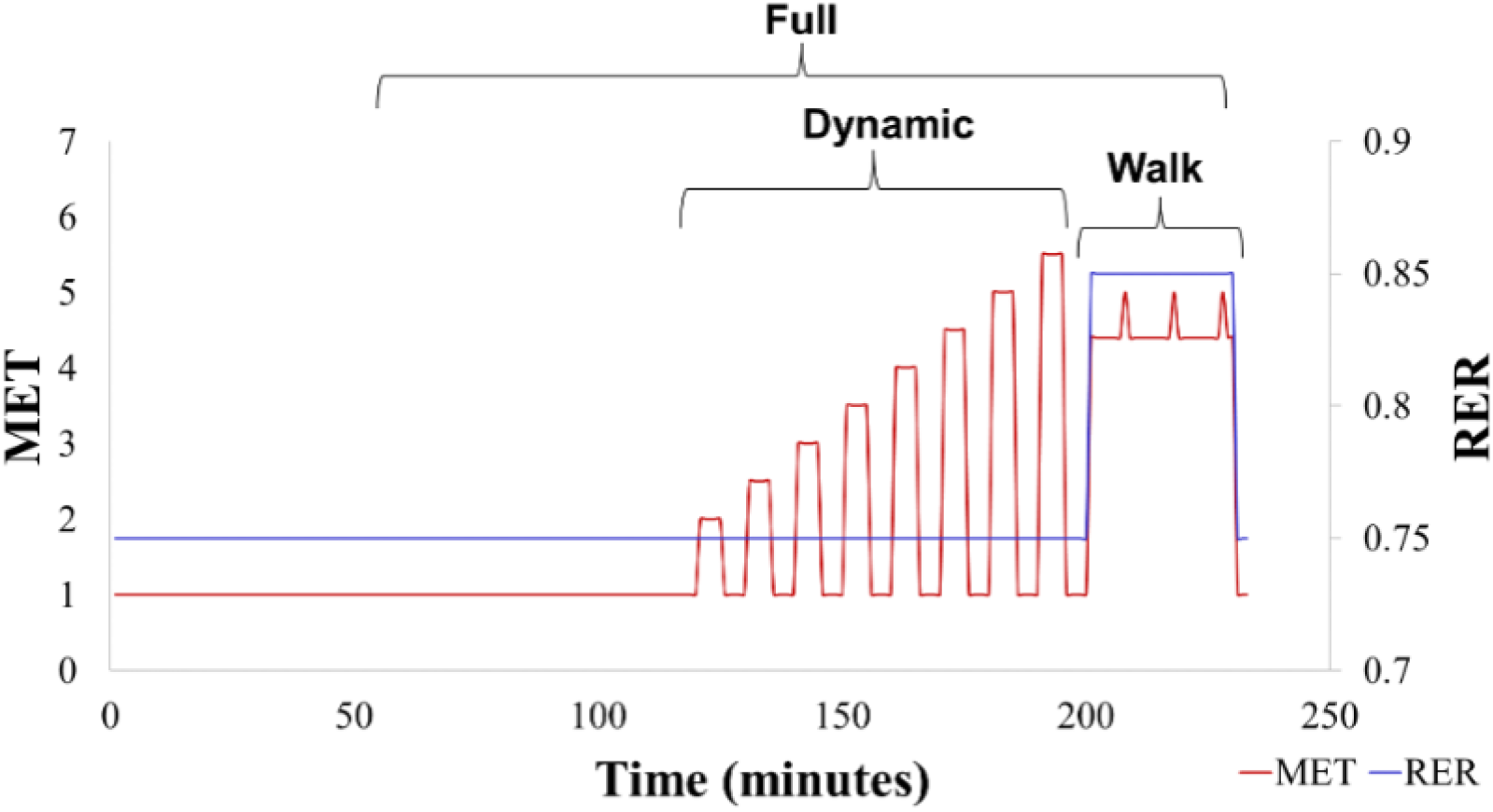
Simulation infusion profile. The infusion protocol began with 2-hour of simulated rest (1 MET, RER 0.75), followed by a dynamic portion with step changes every 5-minute (2-5.5 METS, RER 0.75), and ended with a 30-minute simulated treadmill walk (4.4 METS, RER 0.85) with three ‘challenge periods’ (5.0 METS, RER 0.85).

### 5.2 Calibration of Gas Analyzers

While the hard calibration is an important first step in ensuring accurate measurement, the use of a precision gas blender allows for additional corrections to be made to gas analyzer readings to improve accuracy. We complete “multipoint calibration infusions” that produce twelve O_2_ and eleven CO_2_ concentrations in a steady state stepwise manner (Figure 5. A-B). By infusing these known gas concentrations at different levels, the linearity of analyzer readings can be evaluated across the measurement range with a higher sensitivity than provided by the two-level calibration checks. After infusions, measured vs expected values are plotted and fit with calibration correction equations. These software calibration corrections are then applied to data in post-processing. Shown in Figure 5 C-D is the O_2_ and CO_2_ error across the measurement range from 5 identical multipoint infusions collected over the course of five months. Visual inspection shows that O_2_ readings have a linear offset and that the variation in all three analyzers tends to follow similar patterns within each calibration. Conversely, CO_2_ readings are nonlinear and biased across the measurement range, with each analyzer (grouped by color) expressing a slightly different pattern that appears to be conserved across separate calibrations.

**Figure 5:**
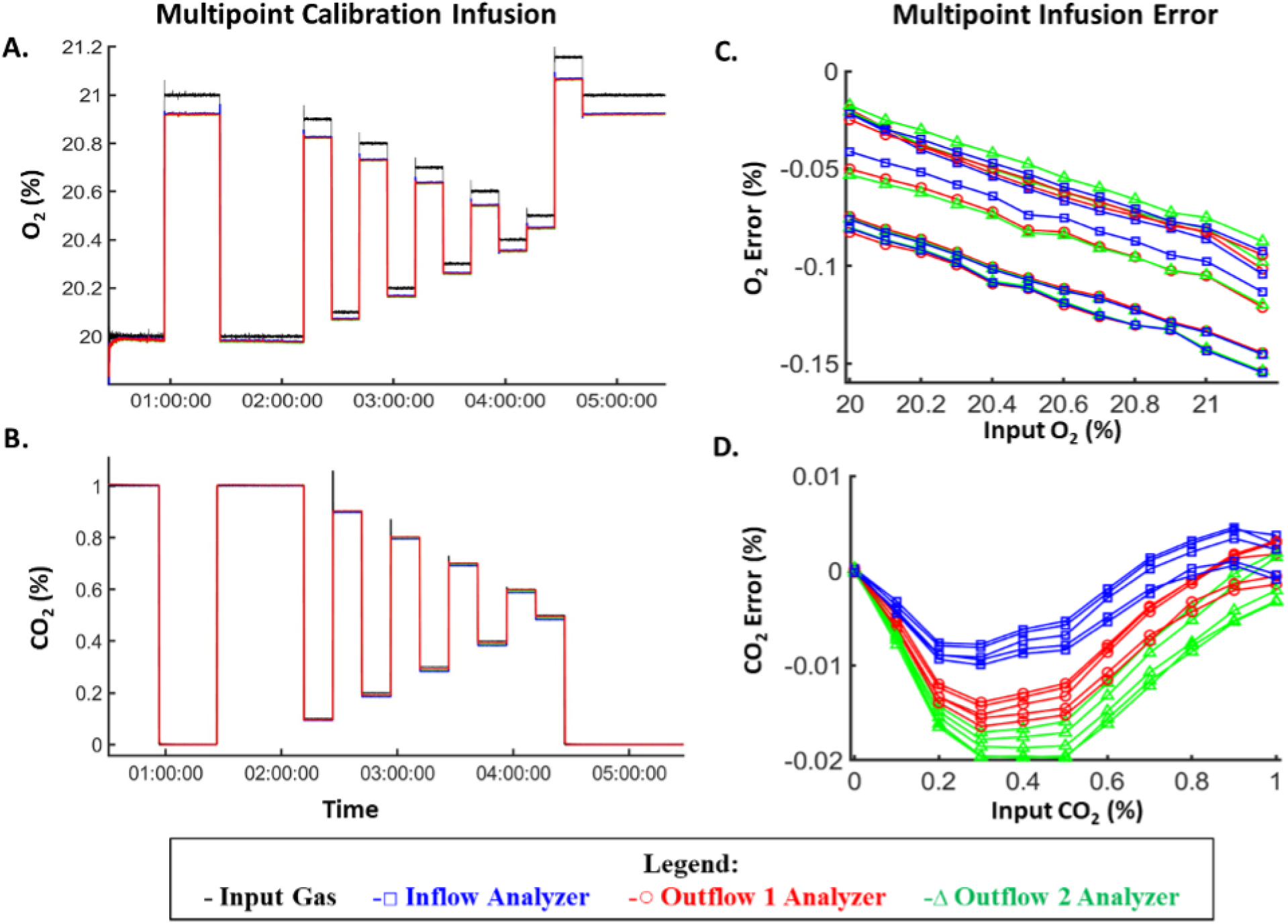
Multi-point calibration infusion data. A-B) O_2_ and CO_2_ expected and measured readings over time during a single multi-point gas infusion. C-D) Variation in O_2_ and CO_2_ readings around linear fits of results from five different multi-point infusions demonstrating the non-linear pattern of CO_2_ measurement error.

To determine the ideal methods for software calibration corrections, we post-processed the simulation infusions described in the previous section with, 1), the two-point hard calibration alone (“no correction”), 2), a linear correction for O_2_ and CO_2_ calculated from our multipoint calibration infusions, and 3), a linear correction for O_2_ and polynomial correction for CO_2_ (Table 1). Applying a linear calibration correction to both O_2_ and CO_2_ resulted in a reversal (underestimate to overestimate) and a very modest reduction in *V̇*O_2_ AUC error and virtually no change in *V̇*CO_2_ error compared to no correction (Table 1). However, applying a linear correction to *V̇* O_2_ and a 3^rd^-order polynomial correction to *V̇* CO_2_ resulted in a considerable reduction in both *V̇* O_2_ and *V̇* CO_2_ error (Table 1). This result supports the use of sensor-specific correction equations and demonstrates how the accuracy of each gas (O_2_ as CO_2_) directly affects both *V̇* O_2_ and *V̇* CO_2_ estimates as suggested in the equation in Figure 1. Considering this observation, O_2_ values are corrected with a linear equation and CO_2_ with a third-order polynomial for the remaining analyses here and in our human research.

**Table 1:** AUC Error (%) by Calibration Correction.

| Correction | Condition | $\dot{V}O_2$ | $\dot{V}CO_2$ |
| --- | --- | --- | --- |
| *No Correction* | Full | -3.5 | -5.3 |
|  | Dynamic | -3.6 | -6.0 |
|  | Walk | -4.9 | -3.4 |
| *Linear $O_2$ & $CO_2$* | Full | 2.9 | -5.4 |
|  | Dynamic | 2.7 | -6.1 |
|  | Walk | 1.6 | -3.6 |
| *Linear $O_2$ & 3<sup>rd</sup> Order $CO_2$* | Full | 1.8 | -0.3 |
|  | Dynamic | 1.5 | -0.2 |
|  | Walk | 0.8 | -0.3 |

Average area under the curve (AUC) error across all 16 simulation infusions processed with 8-minute centered derivatives and various calibration correction equations then segmented into full, dynamic, and walk portions. $\dot{V}O_2$ , oxygen consumption rate; $\dot{V}CO_2$ , carbon dioxide production rate.

### 5.3 Time Derivative

As discussed above, derivative terms are used to reduce noise in the ΔOutflow O_2_% and ΔOutflow CO_2_% terms in the gas exchange equations (Figure. 1) and to account for gas mixing time in the chamber. Thus, instead of calculating the point-to-point change (Δ), the rate of change of O_2_ and CO_2_ in the chamber is averaged over several minutes, which reduces noise but also dampens chamber reactivity. How the selection of different derivative sizes affects measurement error, however, has not been explored. To determine the effect of manipulating the derivative term on the accuracy of *V̇*O_2_ and *V̇*CO_2_, we processed our simulation infusions with 0.5, 1, 2, 4, and 8-minute center derivatives. We then evaluated AUC error for the entire infusion and during the “Dynamic” and “Walk” sub-periods (Figure 4). Because broadening of temporal resolution is likely to have greater effects on shorter epochs, we also evaluated error for the top of each step in the Dynamic sub-period (each 5 min in length). Our results indicate that AUC error is practically unaffected by derivative size when evaluating longer periods of interest (Table 2). However, if the period of interest is shorter, such as in response to a 5-minute activity bout, then greater accuracy is seen with smaller derivative sizes (Table 2). Thus, this data suggests no drawback to the use of derivatives down to 30 seconds in length.

**Table 2:** Average AUC Error (%) by Derivative Size.

| Derv. Size | Full |  | Dynamic |  | Walk |  | Top of Steps |  |
| --- | --- | --- | --- | --- | --- | --- | --- | --- |
| | $\dot{V}O_2$ | $\dot{V}CO_2$ | $\dot{V}O_2$ | $\dot{V}CO_2$ | $\dot{V}O_2$ | $\dot{V}CO_2$ | $\dot{V}O_2$ | $\dot{V}CO_2$ |
| **30 s** | 1.8 | -0.3 | 1.4 | -0.2 | 0.8 | -0.3 | 2.2 | 1.0 |
| **1 min** | 1.8 | -0.3 | 1.4 | -0.2 | 0.8 | -0.3 | 2.1 | 1.7 |
| **2 min** | 1.8 | -0.3 | 1.4 | -0.2 | 0.8 | -0.3 | 3.8 | 4.4 |
| **4 min** | 1.8 | -0.3 | 1.5 | -0.2 | 0.8 | -0.3 | 8.8 | 9.9 |
| **8 min** | 1.8 | -0.3 | 1.5 | -0.2 | 0.8 | -0.3 | 19.4 | 20.6 |

Average area under the curve (AUC) error across all 16 simulation infusions processed with various derivative sizes and then segmented into full, dynamic, and walk portions as well as an average of the top of all five min steps during the dynamic portion.
$\dot{V}O_2$ , oxygen consumption rate; $\dot{V}CO_2$ , carbon dioxide production rate.

To determine if shorter derivative lengths are always beneficial, we measured the absolute error for each possible 5-minute epoch across the entire infusion using a 5-minute moving window and 0.5, 1, 2, 4, and 8-minute center derivatives (Fig. 6). We then calculated the mean absolute error, representing the average error an investigator should expect for 5-minute periods of interest, as well as the standard deviation and maximum absolute error, representing the typical variation and greatest potential error for a given window. Because whole-room calorimeters have traditionally been utilized to investigate metabolism over longer epochs (e.g., 24-hour) we also repeated this analysis with 10, and 30-minute moving windows to determine if there is a minimum epoch length that can be accurately measured (Fig. 6). We found that on average, a window of interest that is 5 min long is nearly as accurate as a 30-minute window (<1.5% difference for *V̇*O_2_ and *V̇*CO_2_), but with double the variation. The maximum error also increases considerably when moving from 30-minute to 5-minute windows (by ∼17% *V̇*O_2_ and 10% for *V̇*CO_2_). As the period of interest shortens, changes in derivative size have an increasing impact on error. For all window sizes, the mean absolute error can be reduced to ≤ 4% ± 4% standard deviation, and the lowest amount of error is consistently found with a derivative of 2 minutes or shorter. The low mean absolute error found with 5-minute windows data demonstrates that whole-room calorimetry can be used for investigating shorter periods of interest than previously believed and that longer derivatives do not always lead to more accurate measurements.

**Figure 6:**
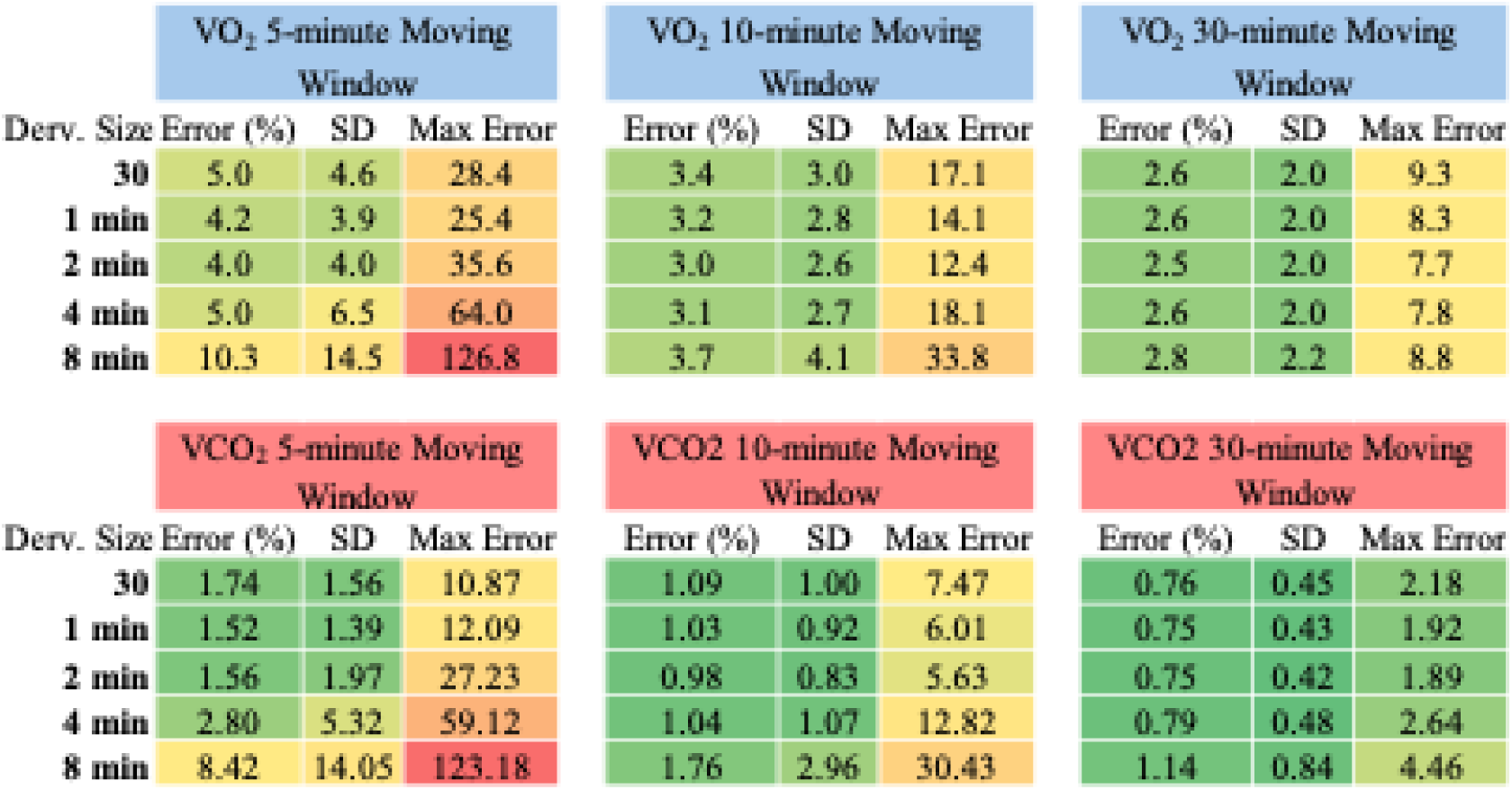
Mean Absolute Error (%) by Period of Interest and Derivative Size. Derivative size, length of the centered derivative used to calculate the rate of change; Error, mean area under the curve error across the course of the infusion; SD, mean standard deviation of the mean error; Max Error, highest error for any single timepoint of the moving window. Results are averaged from 16 simulation infusions ranging from 30-170kg simulated body weight.

If a research question only requires analyzing periods of 30 minutes or longer, then the data in Figure 6 would suggest that derivative size has a negligible effect on error. To determine if this is true, we aggregated the time course absolute error data for the 30-minute moving window and compared the results when either a 30-second or 8-minute derivative was used to calculate *V̇*O_2_ (Fig. 7). As shown in Figure 7, when using either derivative size the absolute error decreases as the signal (oxygen consumption) increases during dynamic steps and simulated walk. However, the use of the 8-minute derivative creates a cyclical error pattern in *V̇*O_2_ estimates that are not washed out by the 30-minute-long epoch. This consistent error pattern would lead to data estimate bias depending on the distance to the nearest transient event (e.g. physical activity). In this case, we can see that despite the greater noise in the 30-second derivative signal there is greater consistency in the signal error. This makes *V̇* O_2_ estimates for a given epoch less susceptible to neighboring events. Collectively, the data presented here suggest that shorter derivative times are likely beneficial to measurement accuracy, even when looking at longer periods of interest (e.g. 30 minutes). However, the shortest derivative does not always elicit the lowest error, particularly with smaller windows of interest (Fig. 6). Therefore, care should be taken when selecting the window and derivative size to balance temporal resolution and potential error based on the hypothesis being tested.

**Figure 7:**
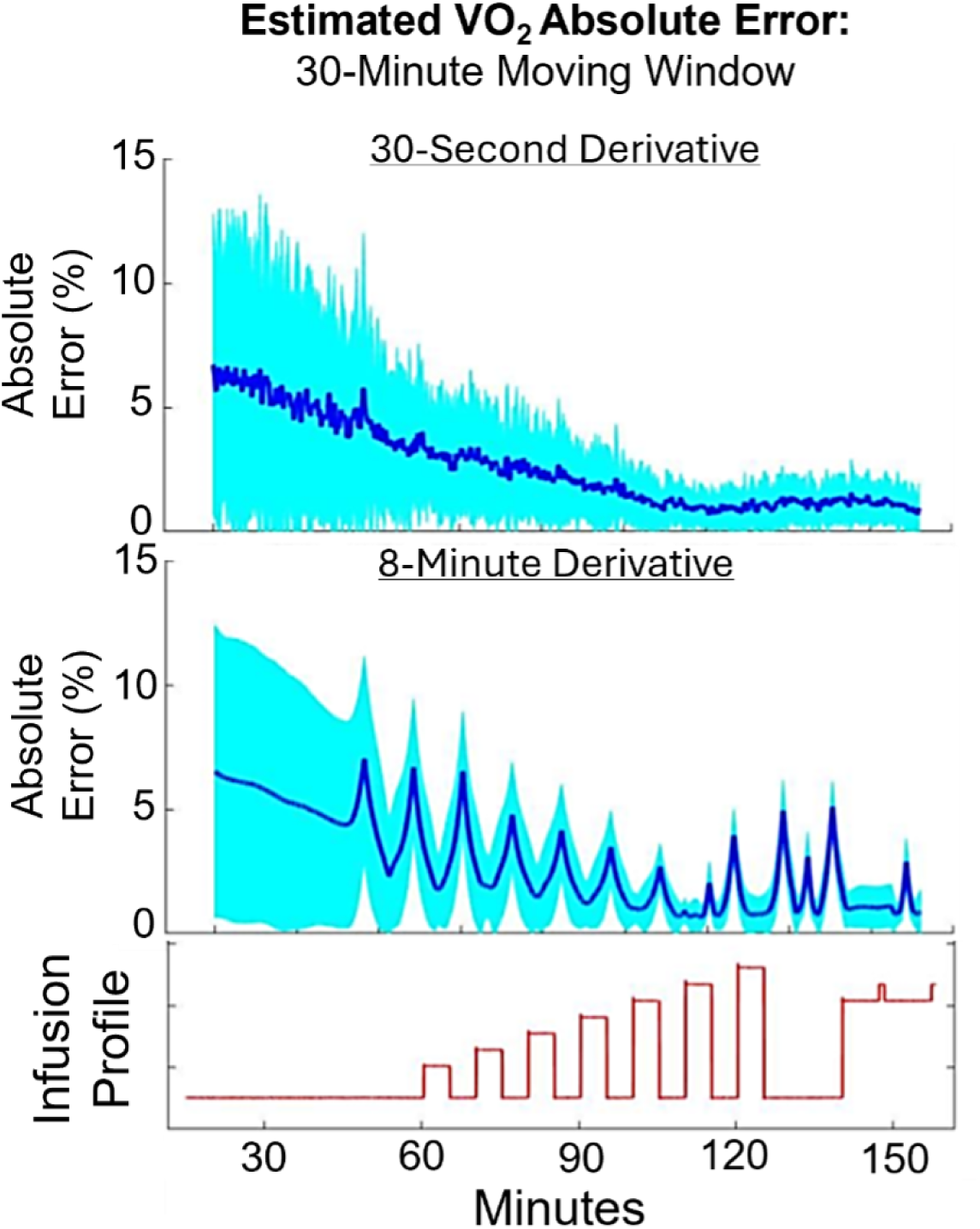
Time Course of Absolute Error in *V̇* O_2_ from Simulation Infusions. Absolute error (dark blue) and standard deviation (cyan) averaged across time from 16 gas infusions. Absolute error was calculated for moving windows of 30 minutes with either a 30-second derivative (top panel) or an 8-minute derivative (middle panel). The expected *V̇*O_2_ pattern is shown in the bottom panel (“Infusion Profile”, unitless).

## 6 Summary

Future investigations of human energy metabolism will be vital to understanding and combatting the current metabolic disease crisis impacting the modern world. Whole-room indirect calorimetry enables metabolic measurement with a balance of accuracy and temporal resolution and has translatability to free-living conditions. We presented methods for calibration, quality assurance, and post-processing procedures to serve as a transparent record for our future work. This includes novel approaches such as a mixed linear (O_2_) and polynomial (CO_2_) calibration correction which reduces error in *V̇*O_2_ and *V̇*CO_2_ considerably. We also demonstrate that the use of smaller derivative sizes improves accuracy when investigating shorter periods of interest and does not decrease accuracy in longer periods. Lastly, we demonstrate that whole-room calorimetry can be used to measure periods of interest as short as 5 minutes with comparable error to 30-minute periods. Our data suggest there is potential to expand the role of room calorimeters to include the impact of shorter duration periods of activity on the traditionally measured 24-hr human energy metabolism. Collectively, we hope this work serves as a reference and building block for improving hardware, data collection, and data processing approaches for the growing field of researchers using such devices.

